# Jointing analysis of scATAC-seq datasets using epiConv

**DOI:** 10.1101/2020.02.13.947242

**Authors:** Li Lin, Liye Zhang

## Abstract

Technical improvement in ATAC-seq makes it possible to profile the chromatin states of single cells at high throughput, but currently no method is available to integrate datasets from multiple sources (different batches of same protocol or multiple experimental protocols). Here we present an algorithm to perform joint analyses on scATAC-seq datasets from multiple sources. In addition to batch correction, we also demonstrate that epiConv is capable of aligning co-assay data (simultaneous profiling of transcriptome and chromatin) onto high-quality ATAC-seq reference or integrating cells in different biological conditions (malignant *vs*. normal), which increases the statistical power of downstream analyses and reveals hidden hierarchy of malignant cells.

## Text

The expression of genes is regulated by transcription factors (TFs) that bind to the regulatory elements of the genome. As the accessible chromatin covers more than 90% TF binding regions, many techniques, such as Assay for Transposase-Accessible Chromatin using sequencing (ATAC-seq), have been developed to detect accessible chromatin^1–3^. Recent technical advancements have made it possible to profile the chromatin states of single cells at a high-throughput manner and along with other molecular modalities (e.g. transcriptome) in the same cells^4–6^. Similar to RNA-seq, ATAC-seq data also suffers from batch effects^7^. But batch correction on scATAC-seq data is more challenging as it is difficult to capture and correct batch derived variations on nearly binary chromatin profiles. For now, there are no specialized batch correction tool designed for scATAC-seq data to our knowledge. Most clustering algorithms for scATAC-seq data capture biological information by learning a series of latent features from binary chromatin profiles. But whether latent features are confounded by batch effects and most importantly, whether batch effects can be removed by scaling these latent features by conventional ways used in scRNA-seq analysis, remain unclear.

Here, we develop a novel algorithm named epiConv (https://github.com/LiLin-biosoft/epiConv) for joint analysis of scATAC-seq data. EpiConv learns the similarities between cells by directly comparing their chromatin profiles (the dot product of two vectors of cells and followed by subsequent library size normalization) (**Fig. 1a**). Comparing to existing algorithms, epiConv performs similarly or slightly better on clustering and embedding (**Fig. S1**). However, its most significant advantage over other methods is that epiConv introduces an additional procedure to remove batch related bias in joint analysis of multiple datasets (**Fig. 1b**): it captures batch derived variations by Eigenvalue Decomposition and remove them by properly scaling the Eigen vectors (See method section for more details). Through comprehensive benchmarking, we demonstrated that epiConv was capable of removing batch effects and in the meantime retaining biological signals.

**Figure 1.**
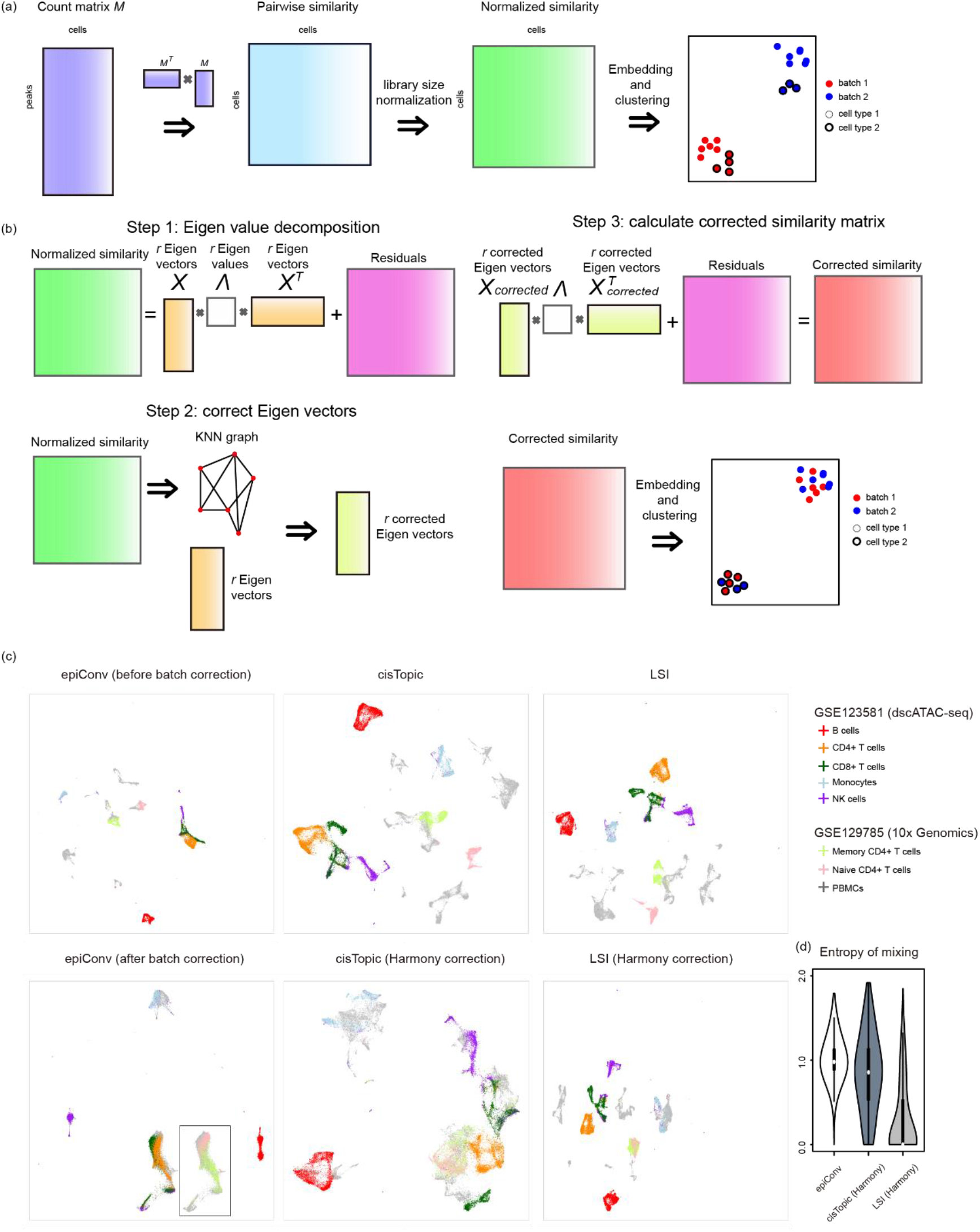
An overview of epiConv algorithm. (**a**) The workflow of calculating similarity matrix between single cells. (**b**) The workflow of batch correction. (**c**) epiConv better corrects batch effects than other methods on a collection of PBMCs datasets. (d) The entropy of batch mixing among cells’ nearest neighbors after batch correction of Figure 1c.

We first applied epiConv to a collection of peripheral blood mononuclear cells (PBMCs) datasets from two studies sequenced by two protocols (10x Genomics and dscATAC-seq) to benchmark its performance of batch correction^5, 8^. For comparison, we also applied two commonly used clustering algorithms of scATAC-seq data (cisTopic^9^ and Latent Sematic Indexing^4^, LSI) to calculate a set of latent features and then removed the batch effects by scaling the latent features using Harmony^10^ (batch correction algorithm developed for scRNA-seq). Without batch correction, all three methods (epiConv, cisTopic and LSI) exhibited obvious batch effects between two studies (**Fig. 1c**). After batch correction, epiConv mixed single cells from two studies together, while cells were still partially or completely clustered by their data source in other methods (**Fig. 1c**). In order to quantitatively evaluate the performance of batch correction, we calculated the information entropy of batch mixing among each cell’s nearest neighbors. EpiConv showed significant improvements of information entropy over other methods, which suggested that epiConv better mixed cells from multiple datasets together (**Fig. 1d**). To evaluate whether cells were grouped based on their biological identities, we performed differential analysis on single cells with known biological identities to get a set of marker chromatin regions for each known cell type. Given that CD8+ T cells grouped into two distinct clusters (naïve CD8+ cluster with accessible *CCR7* promoter and effecter CD8+ cluster with accessible *CCL5* promoter), we performed differential analysis to get markers for both naïve and effector CD8+ clusters. We summed the number of accessible peaks among these markers in single cells to calculate meta signatures that represented their similarities to known cell types. For each cell type, cells with high signature scores from two studies were grouped together and along with the corresponding cells with known identities (compare **Fig. S2** with **Fig 1c**), suggesting that epiConv retained biological heterogeneity.

EpiConv can also increase the resolution of chromatin profiles through joint analysis. Next, we demonstrated that aligning the ATAC-seq profiles of co-assay data (perform scRNA-seq and scATAC-seq simultaneously on the same cell) onto scATAC-seq references overcome the shortcomings of shallow sequencing in co-assay data and improve the performance of clustering and differentially accessible peaks calling. We integrated three datasets together (**Fig. 2a**), one co-assay dataset of mouse adult cerebral cortex^6^ (SNARE-seq), two scATAC-seq datasets of mouse adult whole brain^5, 11^ (sci-ATAC-seq and dscATAC-seq) that served as references, whose sequencing depth are ~5 times deeper than the co-assay data. Single cells from three datasets were mixed together without obvious batch effects by epiConv batch correction (**Fig. 2b**), while other methods fail to do so (**Fig. S3a**). To assess whether the biological heterogeneity was retained after batch correction, we performed conventional clustering on co-assay dataset (**Fig. 2a**) and joint clustering on integrated dataset (**Fig. 2d**, clusters were manually annotated by canonical makers). More than 90% cells from each conventional cluster were assigned to only one joint cluster by epiConv and rare cell types (e.g. C03/Ex 6 and C11/Ex 7, see number of co-assay cells in **Fig. 2d**) were also clearly segregated. On the contrary, other methods were less sensitive to rare cell types recognized by RNA-seq (**Fig. S3b**). We quantitatively evaluated the consistency of clustering by Adjusted Mutual Information (AMI), where higher AMI suggested higher consistency between conventional clustering and joint clustering after batch correction. EpiConv showed the most consistent results among all methods (0.81 in **Fig. 2d**; vs 0.66 and 0.63 in **Fig. S3b**). We also found that conventional cluster C09 could be further grouped into 3 cell types by joint clustering (**Fig. 2c**, clusters highlighted by red circles), which referred to one major cell type (oligodendrocyte) and two rare cell types (oligodendrocyte progenitor cell and microglia) based on their expression profiles from co-assay data (**Fig. 2d**), suggesting that joint analysis also revealed hidden rare cell types. Next, we examined whether single cells of co-assay data were correctly assigned to the reference. In each dataset, we performed differential analysis to get a set of marker peaks for each joint cluster to see whether these markers were conserved across different datasets. Although markers detected from two scATAC-seq references were consistent with each other, the results of co-assay data did not agree with them (**Fig. S4a**). Only markers of big clusters were conserved across three datasets (e.g. Ex 1-4). Moreover, a large number of cluster-specific markers could only be detected from deep sequencing references. We argued that this might be due to shallow sequencing of co-assay data and further incorporate RNA-seq profiles into our evaluation. To do that, we linked each peak to its closest gene promoter in the genome. Then for each cluster, we got a series of associated peaks with its cluster-specific marker genes (**Fig. 2e**). To quantitatively measure the consistency between ATAC-seq and RNA-seq profiles, we calculated the fold change of enrichment for two types of markers (*N_common_* / (*N_ATAC_* * *N_RNA_*) * *N_total_*; *N_common_*: number of shared peaks, *N_ATAC_*, number of chromatin markers, *N_RNA_*, number of marker genes associated peaks, *N_total_*, number of total peaks). We expected that peaks residing near the promoter of one gene were more likely to be its regulators and cluster-specific chromatin markers would enrich in the nearby regions of marker genes if joint analysis correctly clustered recurrent cell types from co-assay and scATAC-seq datasets together. Indeed, we observed the enrichment and for most clusters, much more chromatin markers could be detected from scATAC-seq references than co-assay data and they were with similar or even higher extent of enrichment with marker gene associated peaks (**Fig. S4b**), proving that epiConv correctly assigned co-assay single cells to the references. We noted that there were few chromatin markers of Ex 6 and Ex 7 due to their small cluster size. In order to overcome this problem, we performed the analyses on 7 excitatory neuron clusters only and the results also supported the correctness of epiConv (**Fig. S4c,d**).

**Figure 2.**
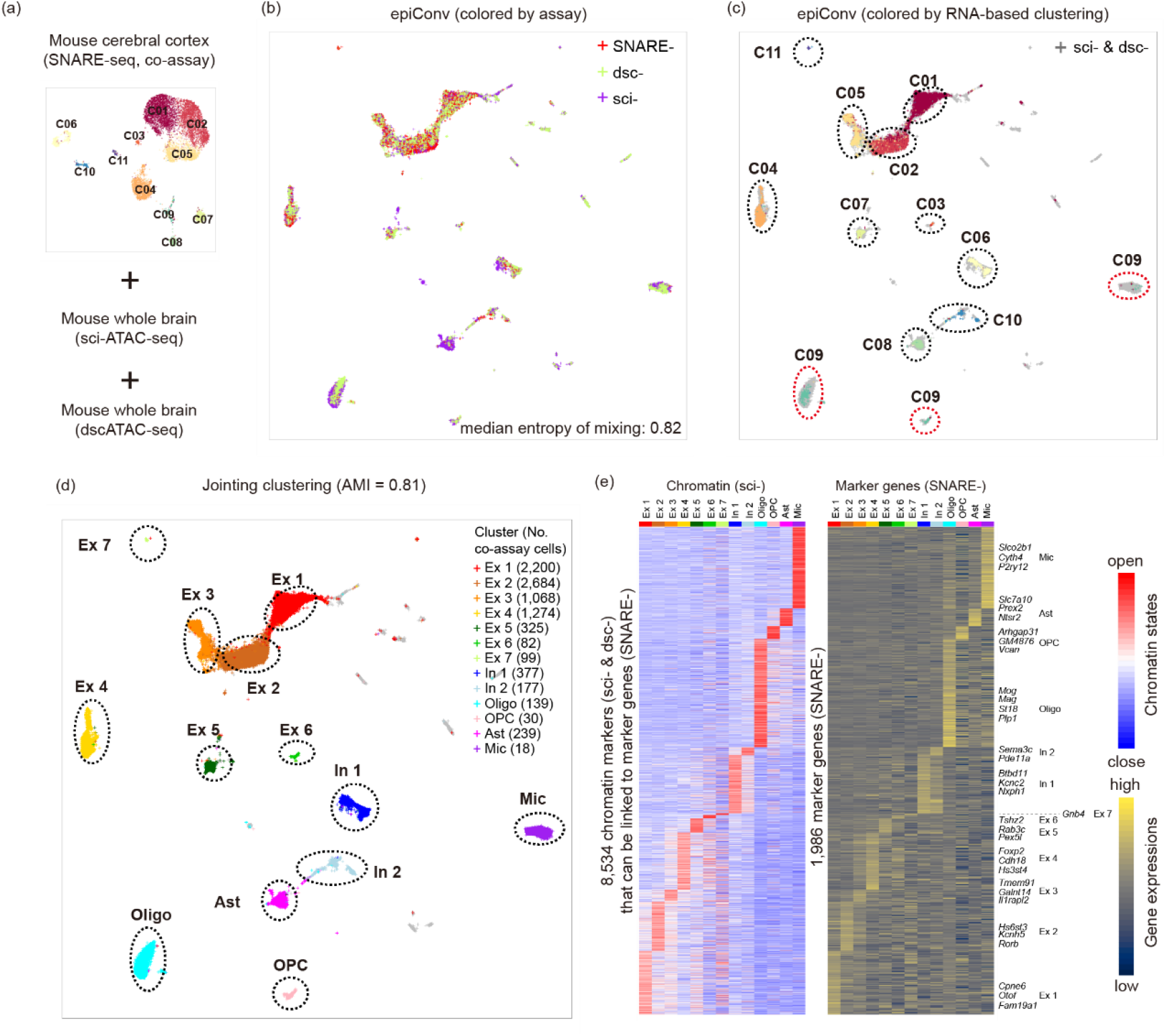
Aligning co-assay (SNARE-seq) data of mouse cerebral cortex to scATAC-seq (sci-ATAC-seq and dscATAC-seq) references increases the resolution of chromatin profiles. (**a**) Low dimensional embedding and clustering of co-assay data. (**b**) Joint embedding of co-assay data and scATAC-seq references, cells are colored by assays. Median value of entropy of batch mixing is shown in bottom right. (**c**) Joint embedding of co-assay data and scATAC-seq references, cells of co-assay data are colored according to conventional clustering in (**a**). Cells of scATAC-seq references are colored in grey. (**d**) Joint embedding of co-assay data and scATAC-seq references, cells are colored by manually annotated joint clusters. Ex, excitatory neuron; In, inhibitory neuron; Oligo, oligodendrocyte; OPC, oligodendrocyte progenitor cell; Ast, astrocyte; Mic, microglia. (**e**) Left: heatmap of chromatin markers detected from scATAC-seq references that can be linked to marker genes. Right: expressions of the corresponding marker genes detected from RNA-seq of co-assay data. Selected marker genes with highest fold changes are shown in the right panel.

We also applied epiConv to another co-assay data of developmental mouse cortex to see whether epiConv performed well on continuous differentiation process. Specifically, we aligned co-assay data of postnatal day 0 mouse cerebral cortex^6^ (SNARE-seq) to scATAC-seq data of E18 cortex, sequenced by 10x Genomics (**Fig. S5a**). By clustering and differential analyses used above, we demonstrated that epiConv correctly assigned co-assay cells to the reference (**Fig. S5b,c**). As the sequencing depth of co-assay data is deeper, we could directly validate the results of epiConv by shared chromatin markers of two datasets. And as expected, we could detect ~5 times more differentially accessible peaks from scATAC-seq reference than co-assay data (**Fig. S5c**). Moreover, we found 2,719 differentially accessible peaks detected from the reference showed consistent pattern with the expressions of their nearest genes along the differentiation path that was examined in original article^6^ (from *Hmgn2^high^* intermediate progenitors to *Cux1^high^* excitatory neurons, J01 to J04). Results based on both distinct cluster and continuous pseudotime assessments proved that epiConv was able to group cells of the same developmental stages together from two independent datasets (**Fig. S5d,e**).

In addition to perform batch correction, integration algorithms could also be used to align cells with different biological conditions or treatments together to identify shared cell populations or perform reference-guided analyses. Here, we further benchmarked the performance of epiConv by aligning malignant cells from mixed-phenotype acute leukemia (MPAL), which was known to present with features of multiple hematopoietic lineages, to normal hematopoiesis^12^. The normal hematopoiesis reference contains bone marrow mononuclear cells (BMMCs) and CD34+ enriched BMMCs and the Leukemia data contains single cells from MPAL patients^12^. After epiConv integration, all malignant cells were projected into several clusters of normal cells (**Fig. 3a,b**). We first annotated clusters by comparing chromatin markers of normal cells with the ATAC-seq profiles of FACS-sorted bulk samples^13^ (**Fig. 3c**; **Fig. S6a**). In addition to previously known cell types, we also found two novel clusters (C08 and C09). The C08 cluster (T biased progenitor) was similar to stem cell cluster (C01) but was more accessible in the marker regions of T cells (~ 50% up-regulated peaks in C08 against C01 are T cell markers) and the C09 cluster (unknown progenitor) was moderately accessible in myeloid, lymphoid and erythroid lineage-specific regions. All malignant samples contained a large proportion of stem-like cells (C01) but the composition of more differentiated cells varied among patients (**Fig. S6b**). The enrichment pattern of cluster-specific markers under normal and malignant hematopoiesis suggested that both abundant and rare leukemic cell types were assigned to the most similar normal cells (**Fig. S6b**). In contrast, malignant and normal cells were still clustered by their biological conditions and patients in the results of cisTopic and LSI (**Fig. S7a,b**). In the original article, Granja *et al*.^12^ developed a novel approach to project leukemic cells to normal hematopoiesis. We found that although it still failed to integrate malignant with normal cells through joint clustering and embedding (**Fig. S7c**), which suggested that the variations between leukemic and normal cells were not corrected, it performed better by projecting malignant cells to an existing embedding of normal cells (add new unseen data points to existing embedding by UMAP), just slightly worse than epiConv (**Fig. 3d,e**). Given malignant cells always contained mixed epigenetic programs of several normal cell types, it was hard to benchmark the results by canonical markers. Instead, we calculated the total number of differentially accessible peaks detected by hypergeometric test under malignant hematopoiesis (higher cluster purity results in more differentially accessible peaks to be detected, which is confirmed by permutation on real data), and the number of recalled markers (markers of one cluster that can be detected from both normal and malignant cells), which reflected consistency of hierarchy between normal and malignant cells. EpiConv showed better performance than Granja *et al*.’s method by detecting more differentially accessible peaks and recalled peaks (**Fig. 3f**). Moreover, malignant cells were not always assigned to the most similar normal cells by Granja *et al*.’s method when comparing the overlap of cluster-specific markers under normal and malignant hematopoiesis (e.g. malignant cells assigned to GMPs by Granja *et al*.’s method were actually more similar to monocytes). Through comparison of cluster-specific markers under normal and malignant hematopoiesis by epiConv, it could be seen that recalled peaks only explained a small fraction of heterogeneity of malignant cells, implying that leukemic cells might have alternative routes to form hierarchy. Moreover, the alternative hierarchy of malignant cells might be easily missed by other methods given that they could not deconvolute highly similar malignant cells into multiple cell types (**Fig. S7**). Motivated by this thinking, we classified cluster-specific markers into three types (**Fig. S8**): 1) hierarchy conserved (shared by normal and malignant hematopoiesis); 2) hierarchy loss (unique to normal hematopoiesis); 3) hierarchy gain (unique to malignant hematopoiesis). The conserved hierarchies were controlled by the transcription factors that were known to be important for hematopoiesis (e.g. RUNX in HSCs and MPPs, PU.1 in myeloid lineages, E2A in lymphoid lineages, GATA in erythroid lineages). While the hierarchies unique to normal hematopoiesis were controlled by the coordination of several epigenetic modifiers, they were largely lost in malignant hematopoiesis (e.g CTCF lost in stem cells and erythroid lineages, C/EBP in myeloid lineages). These results highlighted the power of epiConv to learn the heterogeneity of malignant cells through reference-guided analyses. In this paper, we demonstrate the epiConv performed well in joint analysis of scATAC-seq data under various situations. Moreover, joint analysis provided deeper insights into the epigenetic regulation of single cells of different developmental stages as well as disease conditions. EpiConv is also computationally efficient and can be scaled to datasets with tens of thousands of cells (**Table S1**). We believe that the computational framework in this study along with technical improvements could facilitate better interpretation of the roles of chromatin accessibility in gene regulation in the future.

**Figure 3.**
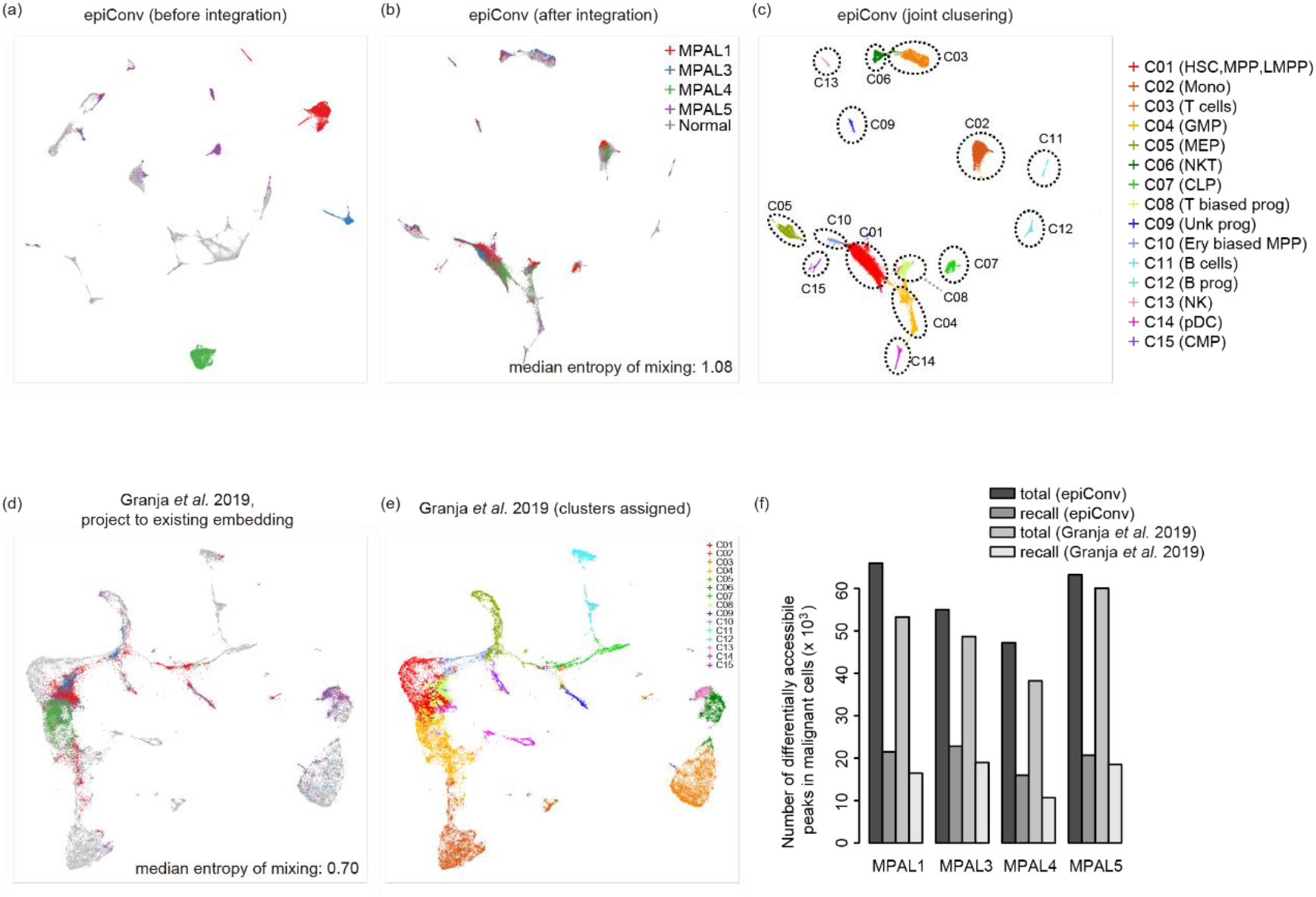
Aligning malignant cells to normal hematopoiesis reveals the hidden epigenetic hierarchy of leukemia. (**a**) Low dimensional embedding of epiConv before integration. (**b**) Low dimensional embedding of epiConv after integration. (**c**) Joint clustering of normal and malignant cells by epiConv. HSC, hematopoietic stem cell; MPP, multipotent progenitors; LMPP lymphoid-primed multipotent progenitor; Mono, monocyte; GMP, granulocyte-macrophage progenitor; MEP, megakaryocyte-erythroid progenitor; CLP, common lymphoid progenitor; Ery biased MPP, erythroid biased MPP; pDC, plasmacytoid dendritic cell; CMP, common myeloid progenitor. (**d**) Low dimensional embedding of Granja *et al*.’s method by projecting malignant cells to existing embedding of normal cells. Cells in (**a**), (**b**) and (**d**) are colored by patients. Median values of entropy of batch mixing in (**b**) and (**d**) are shown in bottom right. (**e**) Cluster assignments by Granja *et al*.’s method. Clusters of normal cells are identical as (**c**) and clusters of malignant cells are assigned based on their nearest normal cells. (**f**) Number of differentially accessible peaks in total and number of recalled differentially accessible peaks under normal and malignant hematopoiesis.

## Methods

### Calculate the similarity matrix by epiConv

EpiConv first calculated the similarity matrix *S* between single cells from a binarized matrix *M*, where rows represent peaks and columns represent cells. The similarity between two cells is calculated by the dot product of two cell vectors, which means that the similarity matrix is calculated by *M^T^M*. After normalization by library size, the similarity matrix can be used for other downstream analyses. A detailed description of this step can be found in Supplementary Note 1.

### Correct batch effects by epiConv

If single cells are from more than one batch, further steps are required to remove the batch effects (**Fig. 1b**). We first apply Eigenvalue Decomposition to normalized similarity matrix *S* and keep *r* Eigen values with largest absolute values and their corresponding Eigen vectors *X*. The similarity matrix *S* can be approximated by matrix product of matrix *X*, which contains Eigen vectors and diagonal matrix *Λ*, whose diagonal elements contains Eigen values. The difference between *S* and its approximation is stored in residual matrix *S_residual_*-Eigen vectors *X* captures batch derived variations and we assume that batch effects can be corrected by properly scaling *X* (**Fig. S9a,b**).

Next, we need to learn the recurrent cell populations across datasets. For simplicity, assume that we have one reference dataset *A* and one target dataset *B* that needs be aligned to the reference (**Fig. S9c**). For each cell in *B*, we find its *k_1_* nearest neighbors in *A* based on the similarity matrix. For each cell in *A*, we find its *k_2_* nearest neighbors in *B*. If one cell in *B* (cell *b)* find a true neighbor in *A* (cell *a*) among its *k_1_* neighbors, we expect that cell *b* should be similar with cell *a*’s *k_2_* neighbors in *B*. To quantitatively measure the similarity between *b* and *a*’s *k_2_* neighbors, we require a collection of features of *B* as guiding features (we will discuss later how to obtain these features). For each guiding feature, we calculate the mean and standard deviation of cell *a*’s *k_2_* neighbors and Z-score normalize the feature value of cell *b* accordingly. If cell *b* shows strong deviation from cell *a*’s *k_2_* neighbors (|*Z score*| > *threshold_Z_*) for any guiding feature, we think cell *a* is a false neighbor of cell *b* and break the link. After such filtering, some cells in *B* may still keep several neighbors in *A* among its initial *k_1_* neighbors (true neighbors) while others may lose most of its initial *k_1_* neighbors. Cells in *B* that have enough true neighbors in *A* (> 5 in this study) are used as anchors for batch correction. For each anchor in *B*, we calculated the difference of Eigen vectors between itself and its true neighbors in *A* as the scaling parameters. The scaling parameters for other non-anchor cells are calculated by the weighted mean of its nearest anchors (10 anchors in this study) on a shared nearest neighbors graph (SNN graph) of *B*, where the weight is equal to the edge of SNN graph. Finally, the corrected eigenvectors *X_corrected_* is calculated by scaling *X* accordingly (**Fig. S9b**).

Parameters settings depend on the complexity and size of data. First, the number of Eigen vectors (*r*) depends on the complexity of data and an appropriate *r* can be easily obtained by manually examining the batch effects on Eigen vectors like **Fig. S9a**. The number of neighbors (*k_1_* and *k_2_*) and threshold_*z*_ control the number of anchors obtained. Smaller *k_1_* and threshold_*z*_ means more stringent filtering and fewer anchors. However, we expected that the number of anchors should be at least ~20% of total cells for each dataset and distributed in all clusters (**Fig. S9d**). As we do not filter cells in *k_2_* neighbors, *k_2_* should be set to a smaller value compared to *k_1_* to prevent from picking too much false neighbors. In this study, we use *r* = 30, *k_1_* = 50, *k_2_* = 10 unless otherwise noted.

Here, we discuss how to obtain the guiding features and SNN graph of dataset *B*. If dataset *B* is scATAC-seq data, the Eigen vectors calculated above can be just used as guiding features given that Eigenvalue Decomposition also captures major biological variations. The SNN graph can be calculated based on the similarity matrix *S* with K-nearest neighbor set to 1% of total cells in *B* (no smaller than 20 or larger than 200). If dataset *B* is co-assay data, we use the Eigen vectors and first 50 principal components (PCs) of gene expressions from RNA-seq as guiding features. The SNN graph is calculated based on the information from RNA-seq or together with ATAC-seq, depending on the data quality of RNA-seq and ATAC-seq (see below for each co-assay dataset). Notably, the guiding features and SNN graph calculated by other methods are also compatible with epiConv if they better resolve the intra-population structure of the dataset.

EpiConv supports of multiple reference datasets and multiple target datasets. When multiple references and target datasets are available, we first integrate all references together (scale Ref *B* to match Ref *A* by finding Ref *B*’s neighbors in Ref *A*, then scale Ref *C* to match Ref *A* and Ref *B* by finding Ref *C*’s neighbors in Ref *A* and Ref *B* …) and then align each target dataset to the references by finding their neighbors in all references.

After the Eigen vectors of all datasets are properly scaled, the corrected similarity matrix is calculated from *X_corrected_*, the diagonal matrix *Λ* and the residual matrix *S_residual_* (**Fig. 1b**).

### Dimension reduction and clustering of epiConv

Given the similarity matrix *S*, we used Uniform Manifold Approximation and Projection^14^ (UMAP) in R package UMAP for dimension reduction and Louvain algorithm implemented in R package Seurat^15^ for clustering. In order to reduce noise, the similarity matrix between cells is first transformed to SNN graph (*S_snn_*) described above. Then, the distance matrix (Max (*S_snn_*) - *S_snn_*) is used as UMAP input. The Louvain clustering is performed on the transformed UMAP graph *G* from the distance matrix (Max (*Ssnn*) - *Ssnn*) to generate consistent results between dimension reduction and clustering. All unmentioned settings are set to default values and the resolution setting in Louvain clustering is manually adjusted between 0.2 and 0.8.

### Detect differentially accessible peaks

We use hypergeometric test to detect cluster-specific accessible peaks. Population size was defined as total number of cells. Sample size was defined as the total library size of cluster divided by the mean library size of all cells. No. of success in the population for each peak was defined as total number of cells with coverage. No. of success in the sample was defined as number of cells with coverage in this cluster. Only peaks that were accessible in at least 1% cells were tested. Peaks with one-tailed p-value smaller than 0.01 were considered as differentially accessible peaks. If peaks were considered as differentially accessible peak in more than one cluster based on the statistical significance threshold, we assigned the peak to the cluster with lowest p-values.

### Peak calling and matrix counting

We used peaks defined by original article or called peaks from ends of fragments by MACS2^16^ (--nomodel --nolambda --keep-dup all --shift −95 --extsize 200) if processed peak file was unavailable. We counted the ends of fragments against peaks to obtain the count matrix. The count matrix was binarized first for further analyses.

### Application of other methods

We applied cisTopic^9^ and Latent Sematic Indexing^4^ (LSI) to scATAC-seq data and used Harmony^10^ to remove batch effects. In cisTopic, the number of topics was set to 20, 30, 40 and 50 and automatically determined. In LSI, we filtered peaks that were accessible in less than 1% cells. The number of reduced dimensions was set to 50. We applied Harmony on the topics learned by cisTopic or PCs learned by LSI without further PCA transformation. Other unmentioned settings were all set to defaults. We performed Louvain clustering on Harmony corrected topics or PCs by Seurat. In order to benchmark batch correction, entropy of batch mixing was calculated by the information entropy of batch among each cell’s 200 nearest neighbors after batch correction, where higher information entropy meant better mixing of cells from different batches.

### Comparison of EpiConv with other methods in individual dataset (without batch effect)

The 5,000 PBMCs data was processed as described above. In order to assess the performance of epiConv and other methods on cell lines data^3^, we clustered cells into 8 clusters (the true number of cell types) by louvain clustering and used Adjust Rand Index (ARI) to evaluate the accuracy.

### scATAC-seq data of PBMCs

We applied epiConv and other methods on two published PBMC datasets. The dataset contains 7 batches of cells. The data from GSE129785 contains two batches of PBMCs, two batches of CD4+CD45RA+ naïve T cells and two batches of CD4+CD45RA-memory T cells^8^. The data from GSE123581 contains isolated CD4+ T cells, CD8+ T cells, CD19+ B cells, CD14+ monocytes and CD56+ NK cells and was treated as one batch^5^. Two replicates of PBMCs of GSE129785 were used as references in epiConv. The threshold_*z*_ was set to 1.5 in this dataset as we found that CD4+ T cells and CD8+ T cells were easily mixed and applied a more stringent threshold. Differential analyses were performed on cells with known cell types. Given T cells shared many common features, we performed differential analyses on CD4+ naïve vs. CD4+ memory T cells of GSE129785 and CD8+ naïve, CD8+ effector and CD4+ T cells of GSE123581 to learn their marker regions instead of comparing them with all other cells.

### scATAC-seq data of adult mouse brain

We integrated co-assay data of mouse adult cerebral cortex from GSE126074^6^ (SNARE-seq), scATAC-seq of mouse adult whole brain from Mouse Cell Atlas^11^ (sci-ATAC-seq) and GSE123581^5^ (dscATAC-seq) together. For RNA-seq of co-assay data, we used a common pipeline of Seurat with default settings (find 2,000 most variable genes and obtain 50 PCs from them) to perform dimension reduction, clustering and finding cluster-specific marker genes. Given that the sequencing depth of RNA-seq is deep but that of ATAC-seq is shallow, we calculated the SNN graph from the first 50 PCs of RNA-seq data only. For ATAC-seq analyses, we first aligned dscATAC-seq data to sciATAC-seq data and then aligned co-assay data to them. The number of Eigen vectors (*r*) was set to 40 and *k_1_* was set to 100 as the reference datasets were large. Dimension reduction and joint clustering were performed on integrated dataset as described above. Performances of clustering by different methods were evaluated by Adjusted Mutual Index (AMI). Given that the resolution of Louvain clustering affected AMI, we performed clustering with resolution from 0.2 to 2.0 for cisTopic and LSI and the highest AMI were reported. For epiConv, we reported the AMI of joint clustering in **Fig. 2d**. The cluster-specific markers shared by sci-ATAC-seq and dscATAC-seq were shown and used for infer the relationships between ATAC-seq and RNA-seq. We also performed differential analysis on excitatory neuron cells instead of all cells to better detect the cluster-specific markers of rare cell types (Ex 6 and Ex 7).

### scATAC-seq data of developmental mouse brain

we aligned the SNARE-seq of postnatal day 0 mouse cerebral cortex^6^ to scATAC-seq data of E18 cortex, sequenced by 10x Genomics. As the sequencing depth of RNA-seq is shallow and we cannot clearly resolve the relationships of single cells based on RNA-seq only, we combined the first 50 PCs of RNA-seq and first 50 Eigen vectors of ATAC-seq to perform low dimensional embedding and clustering using the pipeline of Seurat. The SNN graph used in batch correction was calculated on 100 combined features from RNA-seq and ATAC-seq. Clustering and differential analysis were performed as described above. Clusters of astrocytes (J12) and *Hmgn^high^* Intermediate progenitors (J01) were similar with each other and formed one cluster in unsupervised clustering. We performed further clustering to segregate them. Clusters J14, J15 and J16 were excluded in differential analysis as they did not contain co-assay cells or were mixtures of rare cell types due to lack of proper reference cells. Cell types were labeled based on canonical markers. To investigate the changes of chromatin states and gene expression along differentiation, we also perform differential analyses on ATAC-seq and RNA-seq on 4 clusters (J01 to J04) and selected consistent chromatin markers and gene markers (up-regulated in the same cluster). For cells sequenced by 10x Genomics from J01 to J04, we constructed their pseudotime axis by the sum of the number of accessible peaks among J04 markers minus that of J01 followed by library size normalization as we observed that from J01 to J04, most J01 markers gradually lost accessibility while J04 markers gained accessibility. Next, we imputed the ATAC-seq counts from their 20 nearest neighbors from 10x Genomics data and imputed the RNA-seq counts from their 20 nearest neighbors from SNARE-seq data. Finally, the ATAC-seq counts and RNA-seq counts were normalized by library size with RNA-seq counts further log2 transformed, and they were smoothed along the pseudotime axis by smooth.spline() function in R.

### scATAC-seq data of Leukemia

EpiConv integration, dimension reduction, clustering and differential analysis were performed as described above. There were other two samples (MPAL2 and MPAL5 relapse) in Granja *et al*. 2019^12^. Although we could align them to normal hematopoiesis, they share few common markers with normal cells compared to other samples. We thought it was inappropriate to compare them with normal hematopoiesis and did not show the results of these two samples. To test Granja *et al*.’s method, we first performed LSI on normal cells with 25,000 differentially accessible peaks of lowest *p*-values, kept first 25 PCs, and calculated the PCs of malignant cells using the parameters inferred normal cells. Then we applied UMAP to the PCs of normal and malignant cells (joint embedding) or first applied UMAP to the PCs of normal cells and then calculated the embedding of malignant cells by adding new unseen data points (malignant cells) into existing embedding (projection to existing embedding). We used the clusters from epiConv to annotate normal cells and the labels of malignant cells were assigned based on its nearest normal cell in the embedding of UMAP.

Motif calling was performed by Homer^17^ on the ± 200 bp regions from peak centers. The length of motif was set to 8, 10 and 12. All ATAC-seq peaks were used as background. Motifs that were in fewer than 10% examined peaks were not shown. In T and NK cell lineages, we always called several motifs with low information content and higher GC content. These motifs did not show high consistency with known motifs and were not shown.

### Data availability

All data analyzed in this article are available in public databases and are summarized in **Table S1**. EpiConv is available in Github (https://github.com/LiLin-biosoft/epiConv).

## Acknowledgements

This project was funded by the National Key Research and Development Program of China (2018YFC1004602), National Natural Science Foundation of China (NSF 31871332) and a startup fund to L.Z. from ShanghaiTech University. We would like to thank Vinay Kartha and Xiaoqi Zheng for useful suggestions. We would like to thank Xiaojing Zhao for testing the reproducibility of the study. We would like to thank Yingdong Zhang on his technical support on the HPC platform of ShanghaiTech University.

## Author contributions

L.L. conceived the study, developed the methods and performed the analyses. L.L. and L.Z. wrote the manuscript. L.Z supervised the study.

## Supplementary Materials

**Figure S1.**
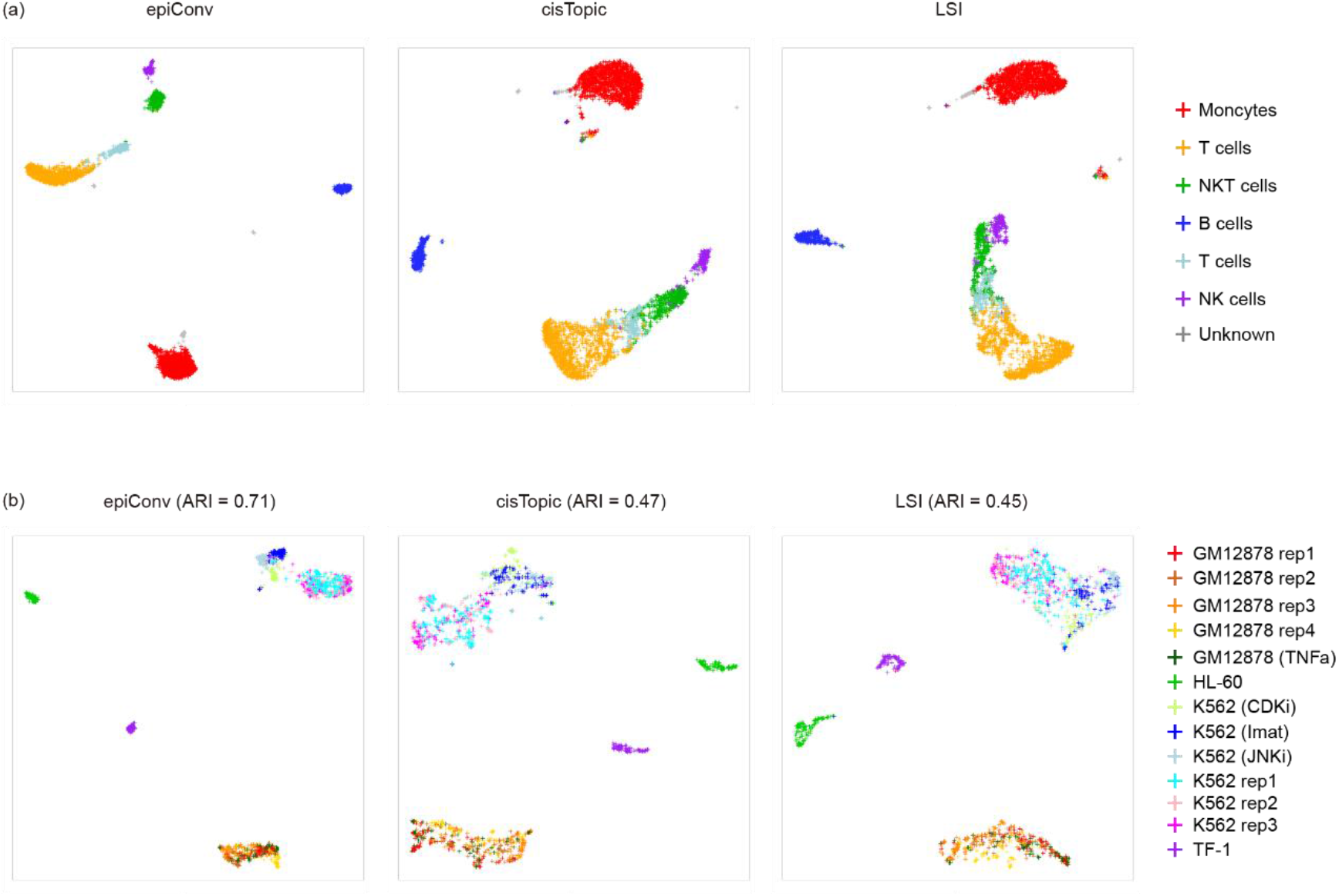
Comparison of epiConv with cisTopic and LSI. (**a**) Low-dimensional embedding of 5,000 PBMCs. Clusters are inferred from epiConv and manually annotated. Results from three methods are largely consistent. (**b**) Low-dimensional embedding of cell lines data. Cells are colored by their identities. Clustering of epiConv achieves higher Adjusted Rand Index (ARI) as it better segregates K562 cells treated by different drugs from untreated cells.

**Figure S2.**
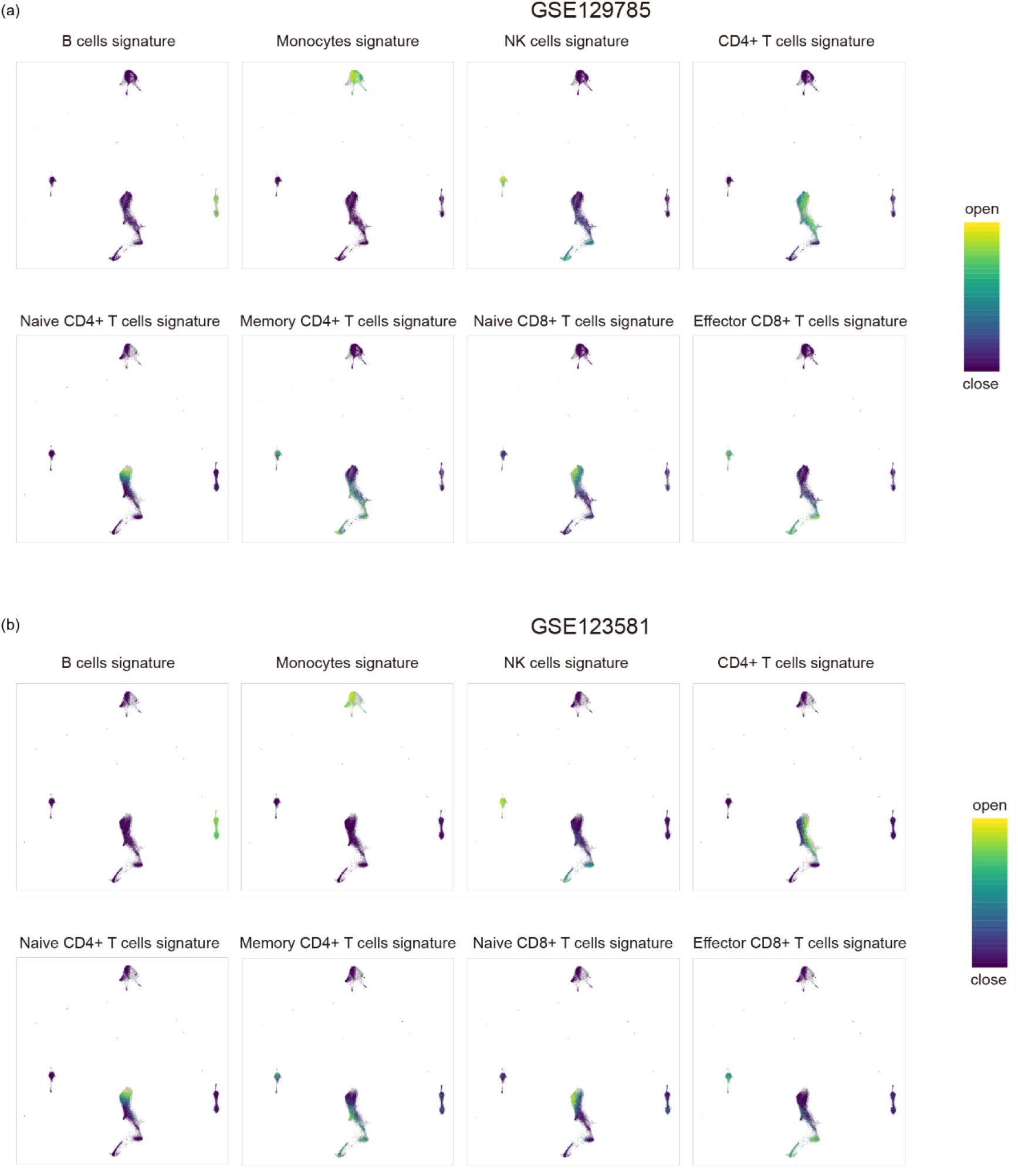
Supplementary figures for **Fig. 1c**. (**a**) Cell type specific signatures of PBMCs from GSE129785. (**b**) Cell type specific signatures of PBMCs from GSE123581.

**Figure S3.**
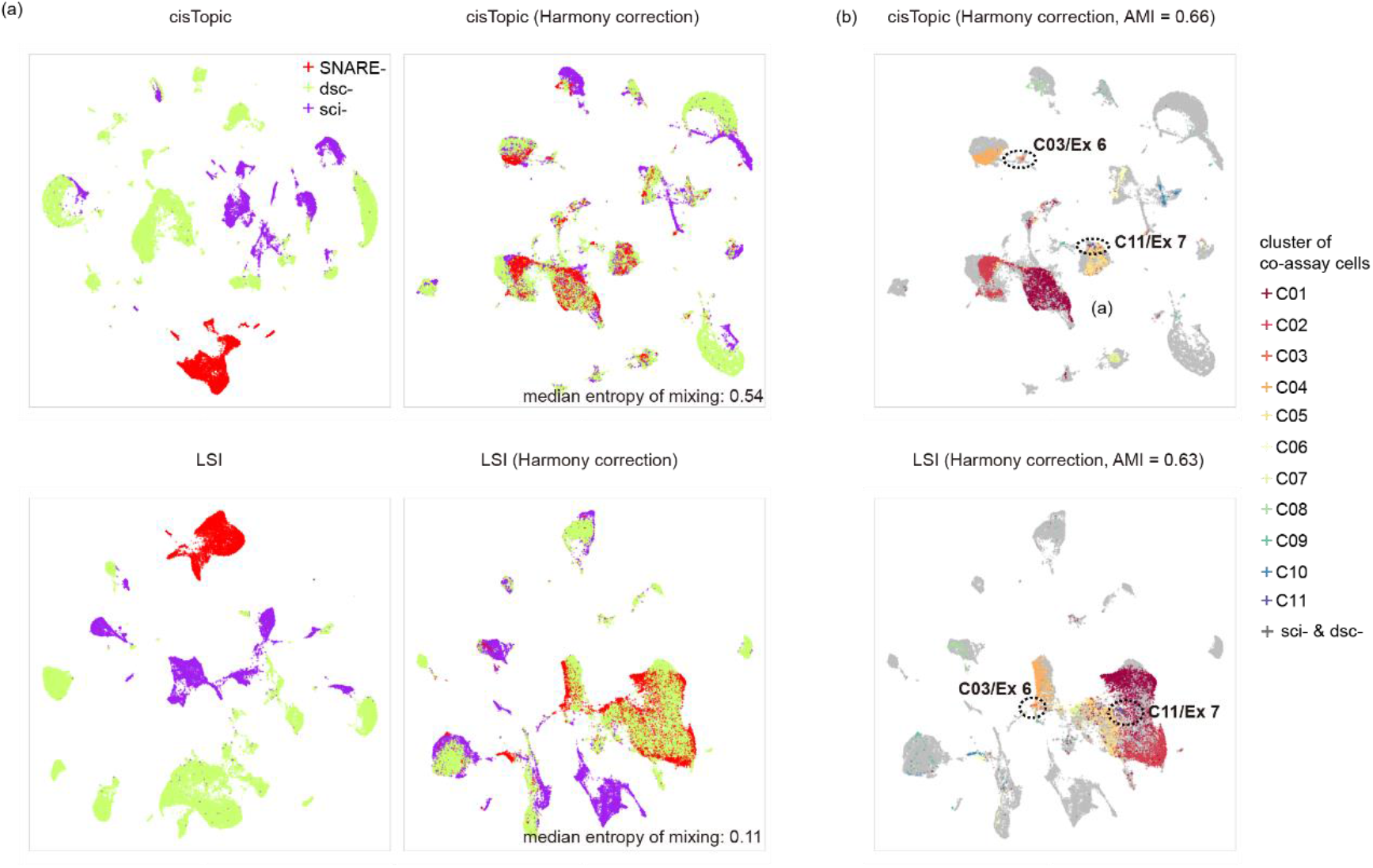
Benchmarking of cisTopic and LSI on adult mouse brain data. (**a**) Low-dimensional embedding of cisTopic and LSI before and after Harmony correction. Cells are colored by assays. Median values of entropy of batch mixing after batch correction are shown in bottom right. (**b**) Low-dimensional embedding of cisTopic and LSI after Harmony correction. Cells are colored according to conventional clustering of co-assay single cells in **Fig. 2a**. Cells of scATAC-seq references are colored in grey. Rare cell types of co-assay data (C03/Ex 6 and C11/Ex 7) are highlighted by dashed circles.

**Figure S4.**
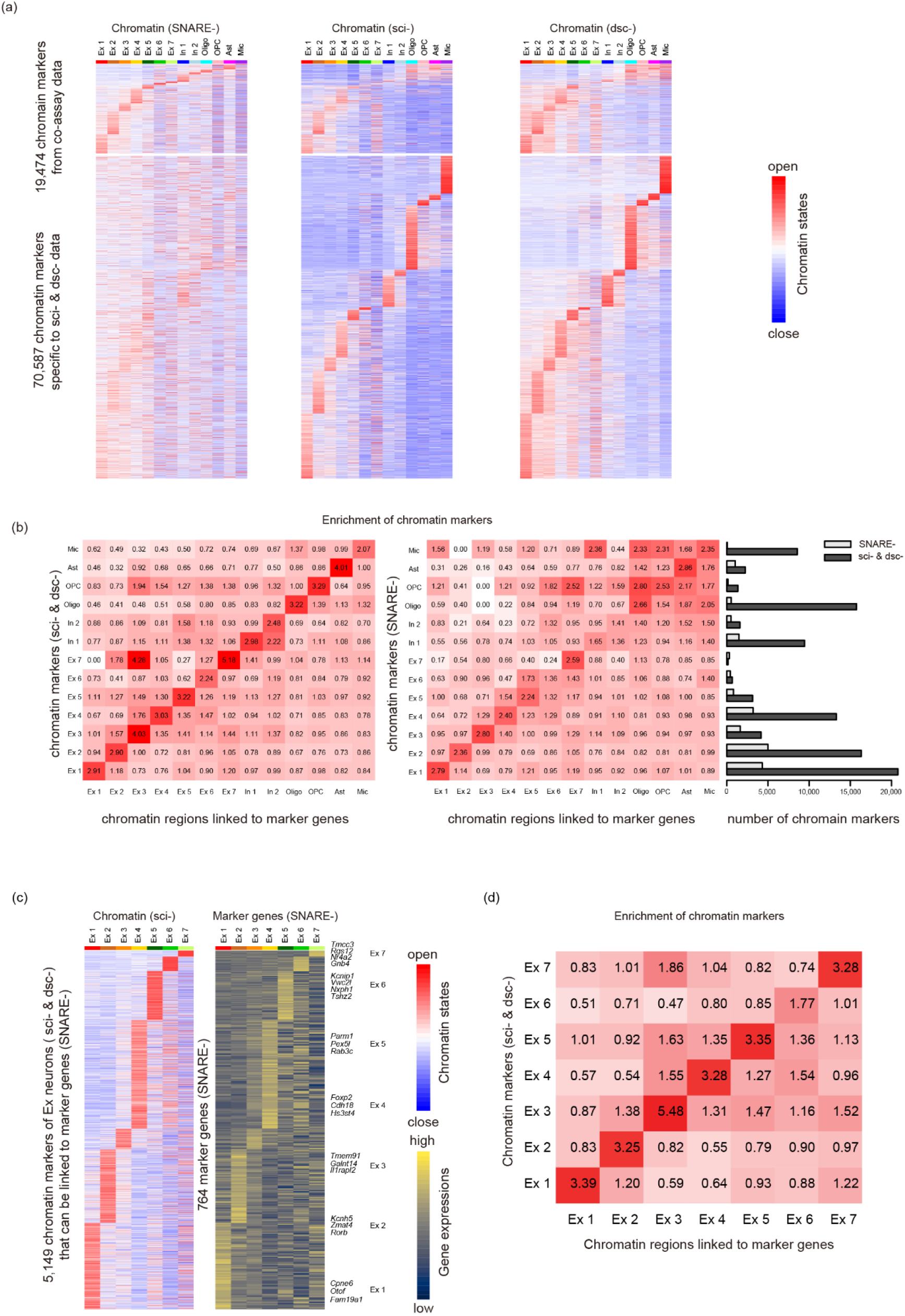
supplementary figures for **Fig. 2**. (**a**) Heatmaps of cluster-specific marker regions detected from ATAC-seq of co-assay data and two scATAC-seq references. Each column represents one aggregated ATAC-seq profiles of cells from corresponding joint clusters. (**b**) Left (scATAC-seq) and middle (co-assay): The fold changes of enrichment between ATAC-seq defined marker regions and RNA-seq defined associated regions. Right: the number of detected markers from scATAC-seq references and co-assay data for each cluster. (**c**) Heatmaps of chromatin marker regions of excitatory neurons detected from scATAC-seq references (left) and the expressions of corresponding marker genes (right). Selected marker genes with highest fold changes are shown in the right panel. (**d**) The fold changes of enrichment between ATAC-seq defined marker regions and RNA-seq defined associated regions among excitatory neurons.

**Figure S5.**
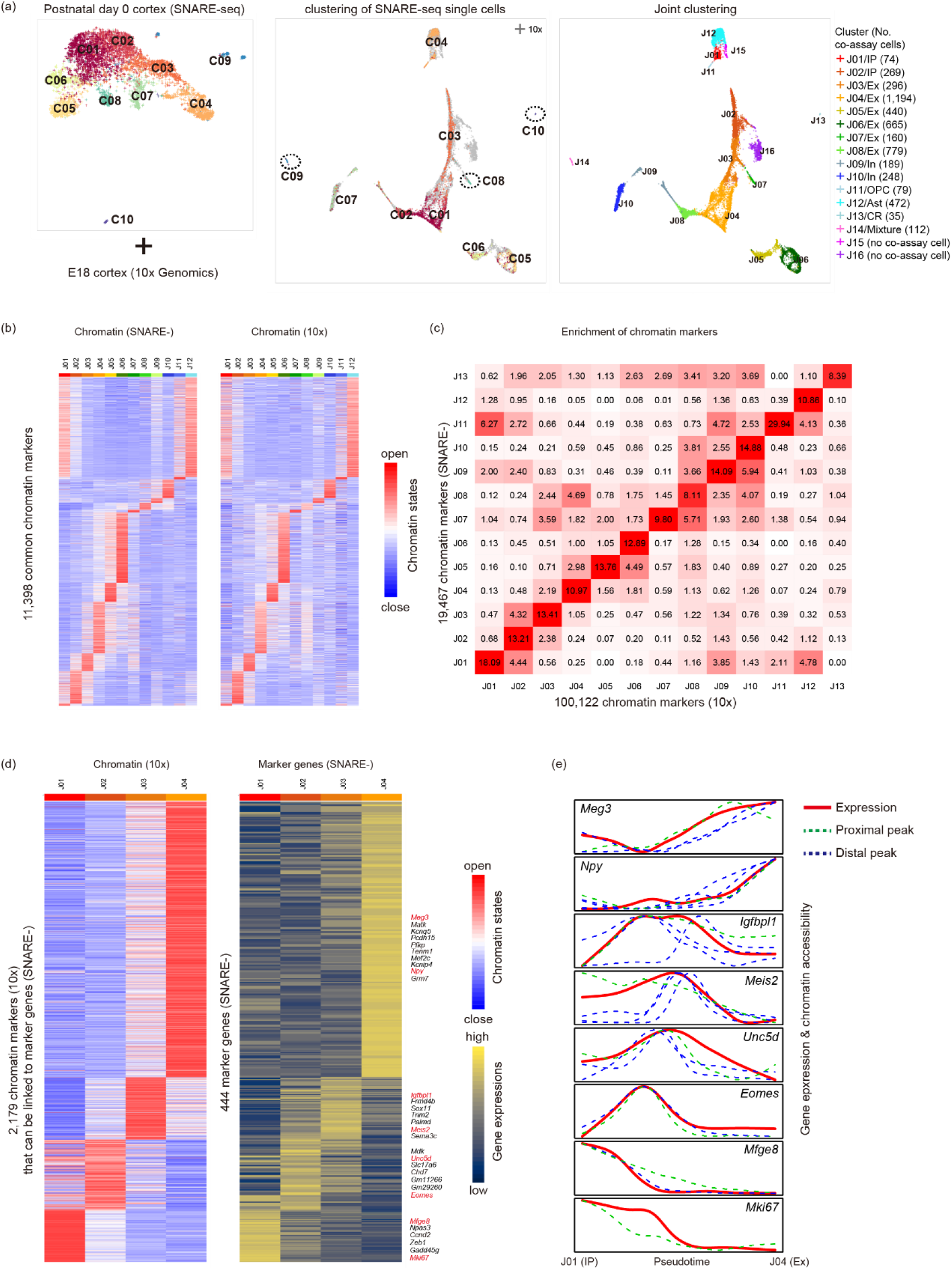
Aligning co-assay (SNARE-seq) data of postnatal day 0 mouse cortex to scATAC-seq (10x Genomics) reference. (**a**) Left: low dimensional embedding and conventional clustering of co-assay data. Middle: joint embedding of co-assay data and scATAC-seq reference, cells of co-assay data are colored according to conventional clustering in the left. Right: joint embedding of co-assay data and scATAC-seq data, cells are colored according to annotated joint clusters. IP, intermediate progenitor; CR, Cajal-retzius cell. (**b**) Heatmaps of cluster-specific chromatin markers shared by two datasets. Each column represents one aggregated ATAC-seq profiles of joint clusters. (**c**) The fold changes of enrichment between chromatin markers detected from ATAC-seq of co-assay data and scATAC-seq reference. (**d**) Changes of chromatin states and gene expressions from intermediate progenitors to excitatory neurons (J01-J04). Left: heatmap of chromatin markers detected from scATAC-seq reference that can be linked to marker genes. Right: expressions of the corresponding marker genes. Selected marker genes with highest fold changes are shown in the right panel. (**e**) Smoothed chromatin and expression profiles of selected makers along the pseudotime of differentiation.

**Figure S6.**
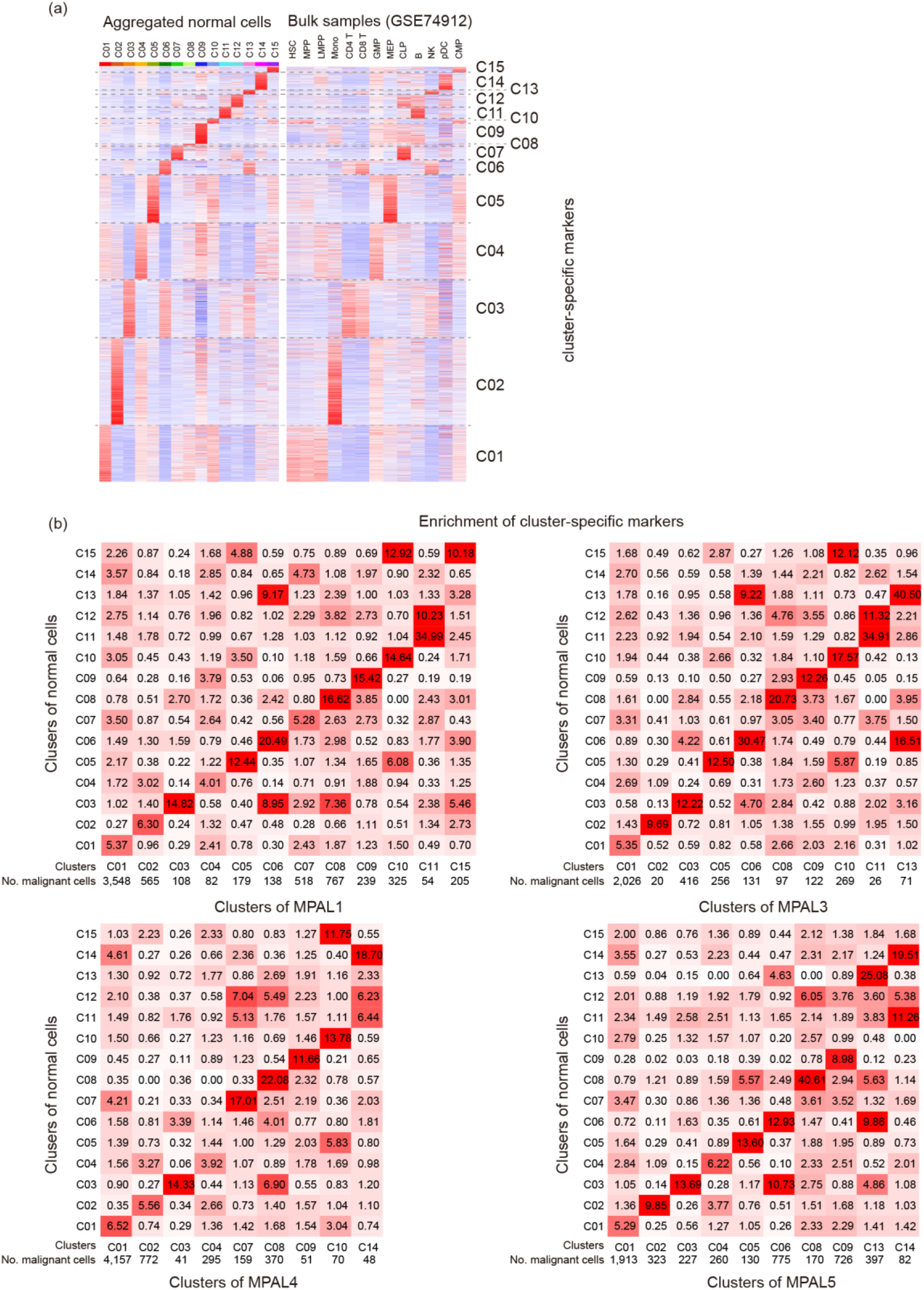
Supplementary figures for **Fig. 3.** (**a**) The aggregated ATAC-seq profiles of normal cells (left) and bulk samples (right) on chromatin marker regions, showing biological identities of each joint cluster. (**b**) The fold change of enrichment between markers detected from normal cells and malignant cells. Number of malignant cells in each cluster is shown on the bottom. Clusters with fewer than 20 cells are not shown.

**Figure S7.**
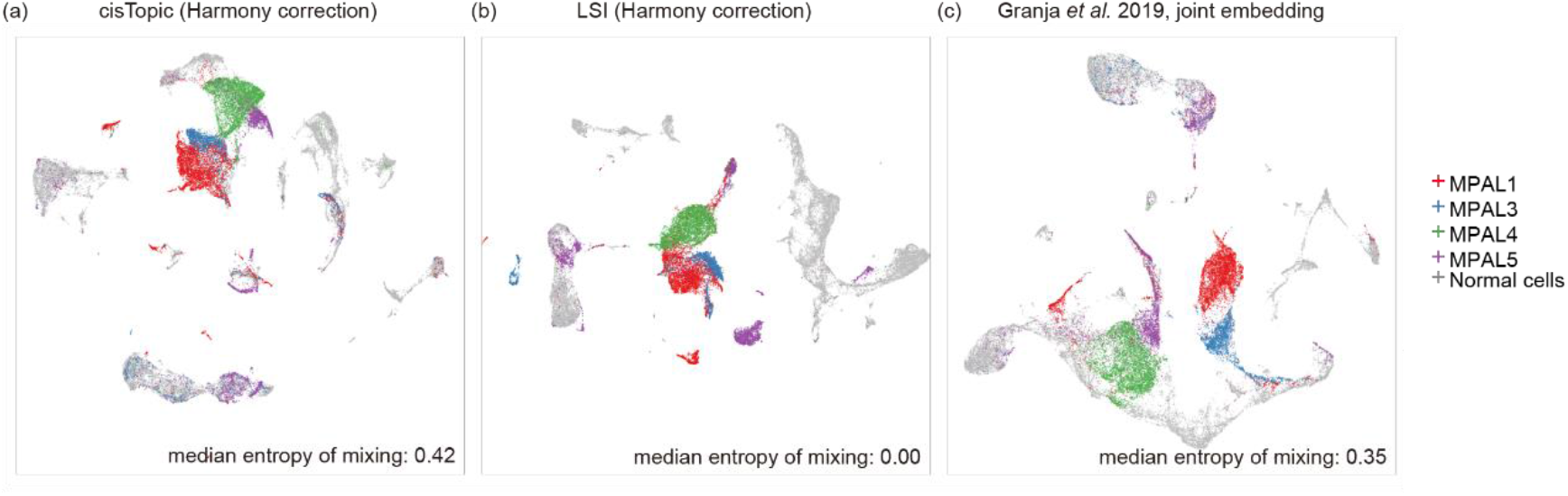
Benchmarking of other methods on Leukemia data. (**a**) CisTopic followed by Harmony correction. (**b**) LSI followed by Harmony correction. (**c**) Granja *et al*.’s method with joint embedding.

**Figure S8.**
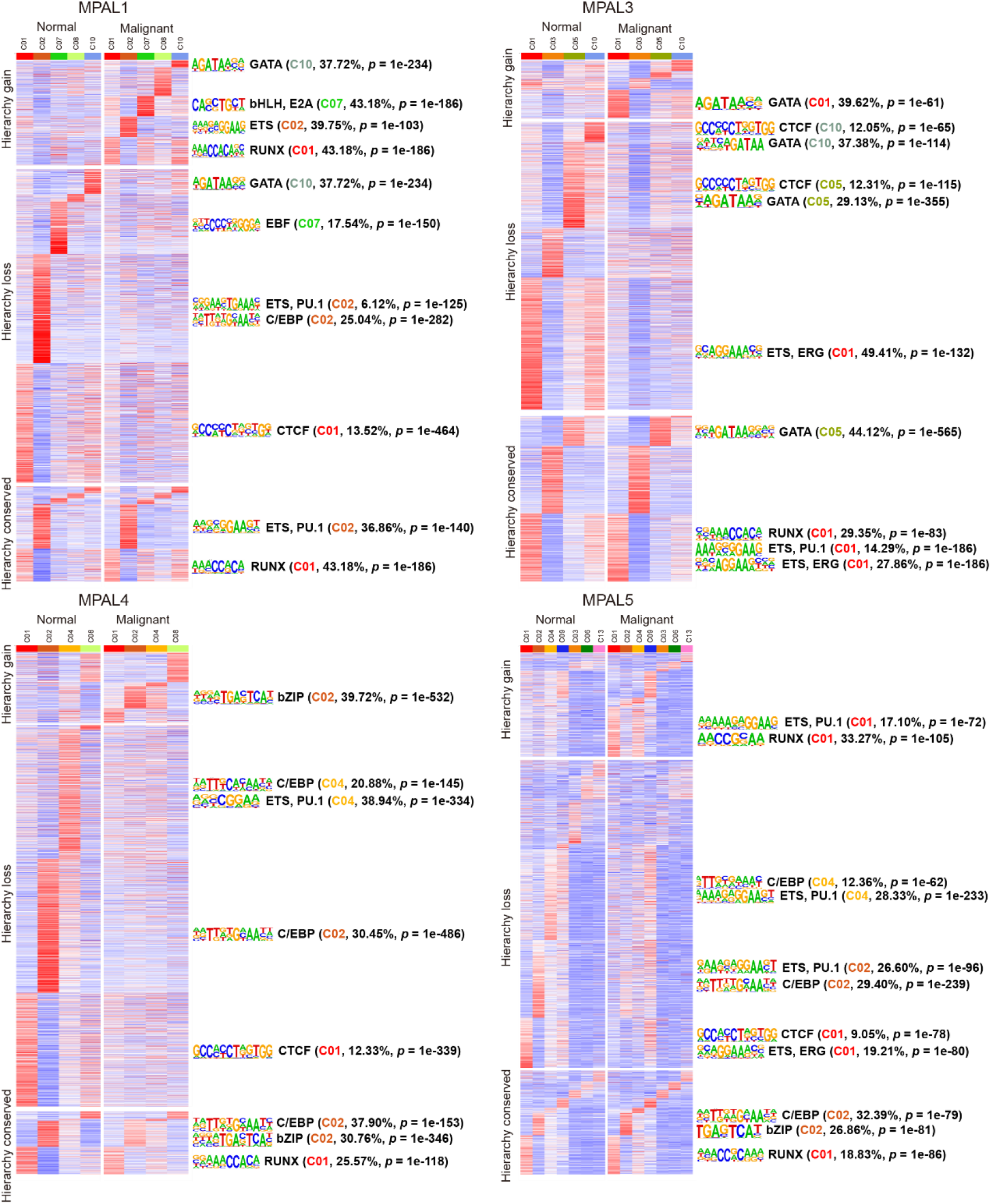
Aggregated ATAC-seq profiles of normal and major malignant clusters on hierarchy conserved, hierarchy loss and hierarchy gain peaks. The enriched motifs with frequency of appearance and *p*-value in each set of peaks are show in right panels.

**Figure S9.**
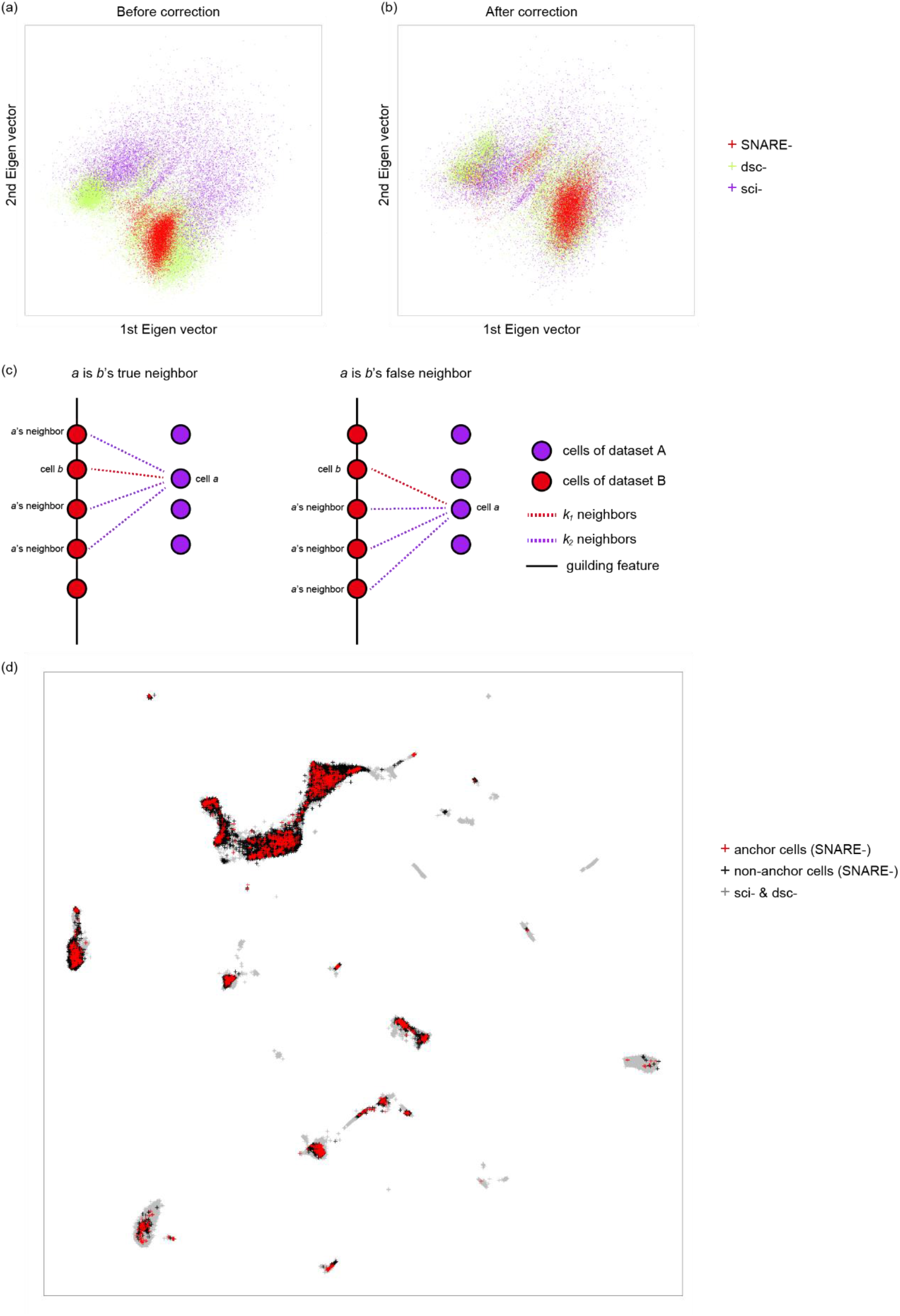
Algorithm details of epiConv batch correction. (**a,b**) The 1^st^ and 2^nd^ Eigen vectors before (**a**) and after (**b**) correction of adult mouse brain data. (**c**) Schematic view of removing false neighbors. (**d**) The distribution of anchor cells from co-assay dataset of adult mouse brain data.

**Table S1.** Published data used in this study with running time and memory requirements.

| dataset | No. cells<br>after QC | Running time<br>(calculate similarity) | Running time<br>(batch correction) | Memory<br>requirement | Source |
| --- | --- | --- | --- | --- | --- |
| PBMCs (**Fig. 1c**) | 41,909 | 5.5 hours | 0.8 hour | < 128 GB | GSE129785<br>GSE123581 |
| Adult mouse<br>brain | 53,762 | 7 hours | 2 hours | < 256 GB | GSE126074<br>GSE123581<br>http://atlas.gs.washington.edu/<br>mouse-atac |
| Developmental<br>mouse brain | 17,613 | 1 hour | 0.5 hour | < 32 GB | GSE126074<br>https://support.10xgenomics.co<br>m/single-cell-atac/datasets |
| Leukemia | 46,510 | 6 hours | 2.1 hours | < 128 GB | GSE139369 |
| 5,000 PBMCs<br>(**Fig. S1b**) | 3,936 | < 5 minutes |  | < 8GB | https://support.10xgenomics.co<br>m/single-cell-atac/datasets |
| Cell lines | 1,174 | < 5 minutes |  | <2 GB | GSE65360 |

## Supplementary Note 1

Here we describe a simple method to calculate the similarities between single cells of ATAC-seq data. First, assume that we have two distributions, *p_x_* and *p_y_*, where cells sample their insertion events from (the chromatin states of two types of cells). If we sample one insertion from *p_x_* and one insertion from *p_y_*, we have a new random variable that is 1 if two insertions occur in the same peak or 0 otherwise. Given random binary vectors *X* with *m* non-zero elements sampled from *p_x_* and *Y* with *n* non-zero elements sampled from *p_y_*, the dot product of *X* and *Y* can be considered as the sum of *m* x *n* variables and is determined by *px* and *py* if *X* and *Y* are sparse. Thus, the *X* · *Y*/(*m* · *n*) can be used to measure the similarity between *X* and *Y*. However, the value of *X* · *Y* is mainly affected by the peaks that have higher frequency. To balance the contribution of peaks with high frequency and low frequency, we weight each peak by *w* = *log*_10_(1 + *inverse frequency of peak*) following LSI, where peaks with higher frequency have lower weight. The observations from real data also support our assumption that the *X* · *Y* can be modeled as *m* x *n* random variables (**Fig. a** below).

In order to reduce the noise of similarity measurement, we adopt a bootstrap strategy. In */*th bootstrap, we calculate *log*_10_ (*X_i_* · *Y_i_*) – *log*_10_*m* – *log*_10_*n*, where we randomly sample some peaks from *X* and *Y* to generate *X_i_* and *Y_i_* (we sampled 20% peaks in each bootstrap and performed 15 bootstraps in this study), and use the mean of *log*_10_ (*X_i_* · *Y_i_*) – *log*_10_*m* – *log*_10_*n* across bootstraps as the similarity between cells.

Even after normalization by *m* x *n*, there are still some dependencies between the similarities and the library sizes. First, similarities between cells with small library sizes tends to have large variations (**Fig. b** below), which is consistent with our initial assumption as the mean value of fewer random variable has larger variation. Second, there are weak negative correlation between the similarity and library size (**Fig. b** below, coefficient ≈ −0.05 in most cases), as the *m* x *n* variables are not independent. We perform linear regression on log transformed similarities and *log*_10_*m* + *log*_10_*n*. The regression residuals are further divided by the corresponding standard deviations, calculated by data points with similar *log*_10_*m* + *log*_10_*n*. The regression residuals after variance stabilization are used as the similarity scores between two cells (**Fig. c** below).

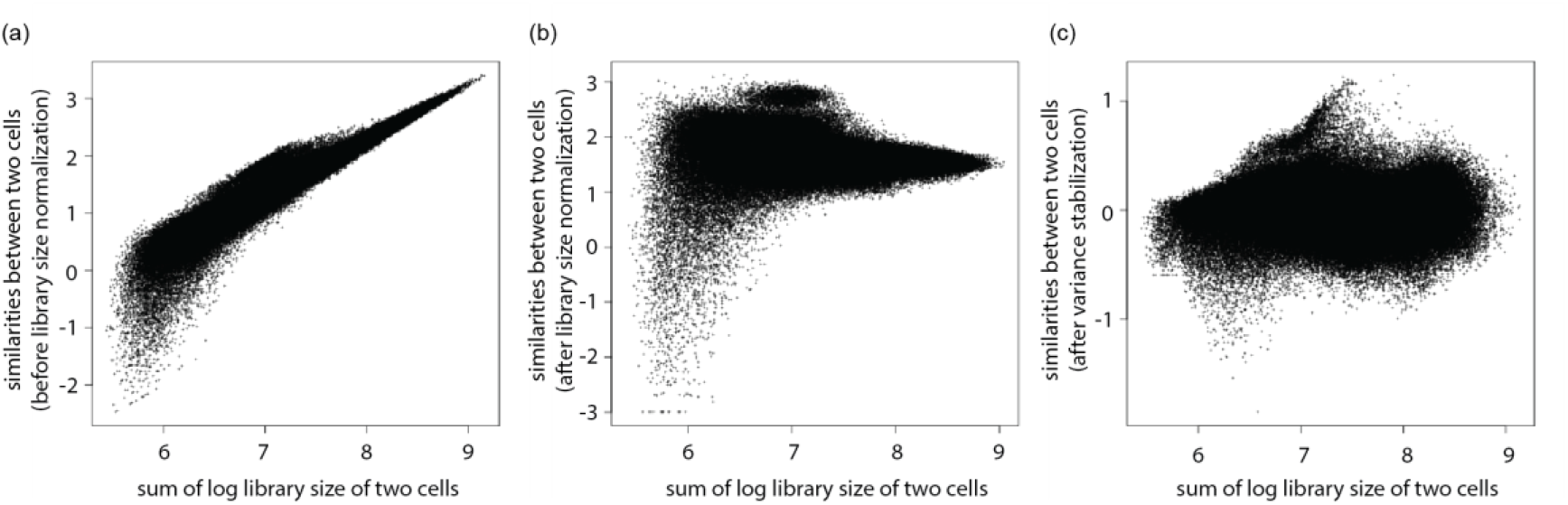

